# Text mining and portal development for gene-specific publications on Alzheimer’s disease and other neurodegenerative diseases

**DOI:** 10.1101/2022.06.28.497987

**Authors:** Jiannan Liu, Huanmei Wu, Daniel H. Robertson, Jie Zhang

## Abstract

**Background:** Tremendous research efforts have been made in the Alzheimer’s disease (AD) field to understand the disease etiology, progression and discover treatments for AD. Many mechanistic hypotheses, therapeutic targets and treatment strategies have been proposed in the last few decades. Reviewing previous work and staying current on this ever-growing body of AD publications is an essential yet difficult task for AD researchers.

**Methods:** In this study, we designed and implemented a natural language processing (NLP) pipeline to extract gene-specific neurodegenerative disease (ND) -focused information from the PubMed database. The collected publication information was filtered and cleaned to construct AD-related gene-specific publication profiles. Six categories of AD-related information are extracted from the processed publication data: publication trend by year, dementia type occurrence, brain region occurrence, mouse model information, keywords occurrence, and co-occurring genes. A user-friendly web portal is then developed using Django framework to provide gene query functions and data visualizations for the generalized and summarized publication information.

**Results:** By implementing the NLP pipeline, we extracted gene-specific ND-related publication information from the abstracts of the publications in the PubMed database. The results are summarized and visualized through an interactive web query portal. Multiple visualization windows display the ND publication trends, mouse models used, dementia types, involved brain regions, keywords to major AD-related biological processes, and co-occurring genes. Direct links to PubMed sites are provided for all recorded publications on the query result page of the web portal.

**Conclusion:** The resulting portal is a valuable tool and data source for quick querying and displaying AD publications tailored to users’ interested research areas and gene targets, which is especially convenient for users without informatic mining skills. Our study will not only keep AD field researchers updated with the progress of AD research, assist them in conducting preliminary examinations efficiently, but also offers additional support for hypothesis generation and validation which will contribute significantly to the communication, dissimilation and progress of AD research.

## Background

Alzheimer’s disease (AD) is one of the most common neurodegenerative diseases. It is estimated that 6.2 million Americans age 65 and older are living with AD in 2021 [1] AD patients suffer from short-term memory difficulties, impaired communications, disorientations and ultimately loss of cognition and fundamental survival skills [2]. AD can be identified by the two hallmark pathologies in the brain, β-amyloid plaque deposition and neurofibrillary tangles of hyperphosphorylated tau [3]. However, the mechanism of how AD initiates and progresses remains unclear. Many AD targets and therapies have been investigated through advanced research and technology, but till now, none has shown significant effect in the general AD population to slow down, stop or reverse the disease progression. Therefore, tremendous efforts have been put into this field for disease mechanism and therapeutical studies.

As a result of these efforts, many scientific publications on AD research have accumulated over the past decades and new research achievements continue to rapidly emerge in the scientific literature. With the wealth of AD-related literature, the AD research community would benefit greatly from an efficiently summarized AD publication mining tool to stay updated with peer research progress, to crosscheck with others results and propose, test and validate their own new hypothesis through the help of such literature mining tool. Such literature mining has helped researchers with hypothesis generation. For example, Malhotra, et al. proposed a disease ontology that specifically focused on AD by using PubMed [4]. Meng, et al. investigated the application of ferulic acid for AD patients by applying text mining methods to PubMed publication records [5]. To further promote the use of the rich research achievements in the AD/ND field and address the urgent need of comprehensive and efficient literature mining, we propose a natural language processing (NLP) pipeline coupled to a web-based neurodegenerative disease (ND) publication mining tool to assist researchers in the AD field. The NLP pipeline extracts the essential ND-related publication from the PubMed database. The web-mining tool offers a convenient means for querying and summarizing related publications for AD-related genes.

## Methods

### Information Processing Workflow

The overall system architecture and information processing workflow are illustrated in Figure 1. We first created a gene list that covers the most commonly studied and well annotated genes in the human genome by using NCBI Gene Database. Next, gene-specific publication profile (GSPP) for each gene is extracted from the abstracts of the publications in the PubMed database using the Entrez module from the *Bio* python package [6]. Once all the GSPPs have been created for genes in the list, the raw data is preprocessed to remove noise and extract the ND-related information from all GSPPs using our NLP pipeline, designed and implemented in six primary modules including publication trend, keywords, dementia types, brain regions involved, mouse models used and co-occurring genes. With the processed data, we then developed a web portal PubAD (https://adexplorer.medicine.iu.edu/pubad/) for users to query the information. The PubAD provides gene query functions and associated visualizations such as bar plots, word cloud plots, lollipop plots, etc., enabling easy exploration and interpretation of our processed data. The detailed information of each step is further illustrated below.

**Figure 1.**
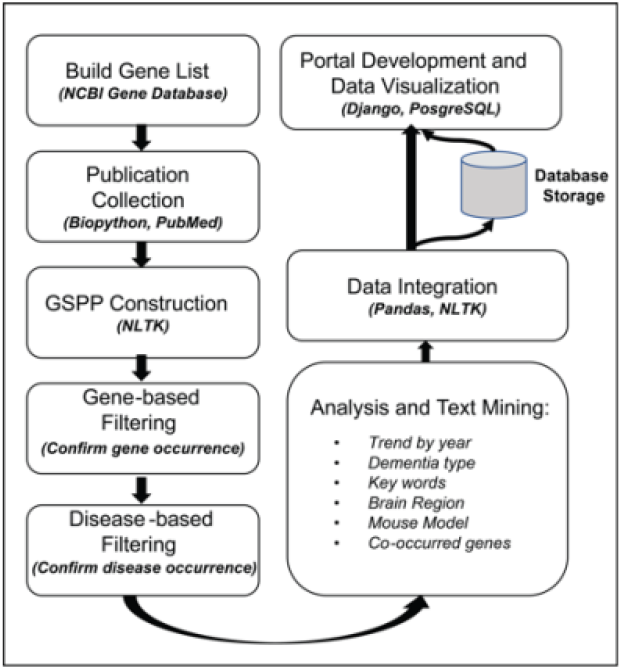
Overview of the PubAD data processing workflow.

### Data Collection

The Entrez module from the Bio python package is used to access the publications and source metadata using the PubMed API [7]. The Entrez module is a python wrapper around the Entrez Programming Utilities (E-utilities) which allows us to search, fetch and parse data from NCBI databases programmatically. The HUGO gene symbol is used as the primary identifier for each gene, while the gene’s aliases are also collected from the NCBI Gene database. To build a GSPP, we extracted the relevant data with a focus on the queried gene’s information appearing in the publication titles and abstracts to reduce the complexity and redundancy of GSPPs[8]. This approach assumes that the gene’s official symbol (or its aliases) occurring in the title or abstract indicates the gene is an important component of the study in that particular publication. For each publication in a GSPP, the publication information was extracted including the title, abstract, year, PMID and journal name.

As the total number of publications in the PubMed database is extremely large, the data collecting process is time-consuming. We therefore implemented a PubMed data extraction pipeline using the Python multiprocessing package to speed up the data collection process by fully leveraging multiple processors on a given machine. Once all the publication information is obtained for each GSPP in JSON format, further data processing is performed to transform each gene ‘ s publications into a single CSV file. Each row is a record of one publication, while the columns are gene-specific publication information mentioned above (year, PMID, etc.).

Upon collecting all GSPP files, we designed and implemented an NLP pipeline to clean and extract the ND-related information from all GSPPs using five primary steps: data cleaning, dementia type detection (the seven major dementia types are defined as AD, Parkinson’s disease, Huntington’s Disease, frontotemporal dementia, Lewy body dementia, vascular dementia and mixed dementia), key ND-related biological processes extraction, publication trend by year, and gene co-occurrence statistics (Figure 1). These steps ensure an accurate summarization and analysis of the essential information for each gene from the ND-related publications, guided by experts with years of experience in AD research to ensure that the extracted categorical information was concise, inclusive, and relevant to the AD.

### Data Cleaning

The official HUGO gene symbol and its aliases are used in data collection to build the GSPP. However, directly querying the PubMed database may result in redundant publications in which the gene name did not appear in either the title or abstract. Therefore, we included an gene filtering step to confirm the gene symbols or their aliases are actually in the title or abstract. Once all GSPPs are obtained, the sentences are tokenized using tokenizer based on Punkt sentence tokenization models[9] to obtain the word tokens in publications title and abstract, all punctuations are removed from the resulting tokens. Finally, current gene’s symbol or aliases are searched against the filtered tokens to check occurrence. Publications that do not have the current gene symbol or its aliases from GSPP are removed.

Additional data cleaning is based on diseases of focus. As one type of NDs, the symptoms and mechanisms of AD are highly related to those of other ND types. AD researchers can be inspired by the results from studies in other types of dementia. Therefore, the diseases of focus in this study include AD, Parkinson’s disease, Huntington’s Disease, frontotemporal dementia, Lewy body dementia, vascular dementia and mixed dementia. To filter for the publications in GSPP related to the diseases of focus, we constructed a vocabular dictionary to include all names of diseases of focus and their semantic variants by using simplified names, for example, “Alzheimer’s Disease” is represented by “alzheimer”, then we applied similar data processing steps as in the gene filtering step using NLTK package to tokenize and clean the data. The resulting tokens are used for examining the occurrence of these diseases in a publication’s title and abstract. The filtered GSPPs are used for further information retrieval.

### Information Extraction & Visualization

For each filtered clean GSPP, information extraction was performed for six primary categories of information: publication trend by year, dementia type occurrence, brain region occurrence, mouse model information, keywords occurrence, and co-occurring genes. For each category, the occurrence count of the corresponding sub-category (such as hippocampus in brain region) was collected together with the publications’ PMIDs for detailed information of publications.

1. *Yearly publication trend*: For each GSPP, the publications are stratified by year, with the publication PMIDs and the total count for any specific year. This yearly information is represented as the Python dictionaries used in subsequent analyses and data mining of gene-related information. For example, one ongoing project we are working on identifying the promising potential biomarkers based on the gene publication trend.
2. *Dementia type occurrence*: To accurately identify the dementia types mentioned in the data cleaning section, we tokenized the paper title and abstract in each GSPP. If a dementia form occurs in the tokens, the publication and its PMID are added to the dementia-specific publication list. Currently, our system only gathers total dementia-specific publications for all years. However, it can be further stratified into yearly publication on a specific dementia type.
3. *Brain region information*: Our study identifies brain region information based on thirteen brain regions commonly studied in neurodegenerative diseases [10, 11]. In alphabetical order, these thirteen brain regions are Amygdala, Basal ganglia, Brain stem, Cerebellum, Cingulate gyrus, Corpus callosum, Hippocampus, Hypothalamus, Neocortex, Pituitary gland, Prefrontal cortex, Spinal cord, and Thalamus. The approach to extract the brain region information is similar to that used for dementia types: Tokenize the title and abstract of a publication and then check if a specific brain region is included in the tokens.
4. *Mouse model information*: AD-related mouse models are valuable for investigating disease pathologies and preclinical testing [12]. There are many publications with novel findings from experiments on AD mouse models. It assists AD researchers to access the published studies carried out with specific AD mouse models. We collected the marked “available” mouse model strain information from the mouse model strain table on the MODEL-AD website [13]. The strain name is used as the primary identifier for the mouse model information extracted from the publication titles and abstracts, similar to dementia types and brain region information extraction.
5. *Keywords occurrence*: Our AD research experts identified 32 representative keywords related to AD research (see Supplement Table-1 for a full list). For example, ‘tau deposition’ and ‘innate immune response’ are in the keywords list because tau deposition is considered as one of the most important characteristics of AD development [14] while ‘innate immune response’ was reported that peripheral immune cells are actively involved in AD progression [15]. With the publication count for each key word, a word cloud plot is generated using *wordcloud* python package[16], the font size of the key word indicates the occurrence frequency of the key word in a GSPP.
6. *Co-occurring genes*: Identifying which genes are commonly mentioned together in one publication empowers researchers to broaden their research scope and potentially discover the related mechanisms by investigating highly related genes in publications of interest. To improve the accuracy of gene name annotation in literature, we performed parts of speech (POS) tagging on tokenized publication title and abstract. Only tokens that have certain tags are used for gene name identification, these tags include NNP, NN, NNS, JJ, etc. Then all genes that occurred in the filtered tokens were annotated with official genes symbols. As mentioned in the data cleaning, each filtered GSPP is marked for a specific gene. The occurrences of other genes will be identified and aggregated to get the co-occurring genes information.

### Backend Database Design

With the established publication library, we developed a website using the Django framework [17] which incorporates a user-friendly interface for publication access and analysis. A PostgreSQL database [18] was designed to manage the collected raw data and processed information. We used a large database table to record six categories of processed information, each category of information has two corresponding attributes, one is for storing the publication count information, another one for storing the related PMIDs. In the processed information table, HUGO gene symbol is served as primary key, all attributes for processed information are set to be JSON type, each row of the table stores all processed information for a certain gene. Detailed gene information such as genome location, aliases, etc. are collected from NCBI Gene database and stored in a database table to serve as reference information.

### Web Portal Development

By using Django which is a python implementation of Model-View-Controller (MVC) framework, we created an automated workflow that can extract processed publication information from the database with user inputs and then generate interactive visualizations (Figure 2). With the count information and PMIDs in the processed data, we designed and implemented a visualization dashboard with D3.js [19] to provide intuitive illustrations for five categories of information. Each category of information is organized in one tab. Each tab of the dashboard provides different visualizations together with a description of the current information category. The layout of the website is structured with Bootstrap 4 [20]. The search box of the home page incorporates the autocomplete function of Ajax [21] to help with real-time gene identification with users’ inputs.

**Figure 2.**
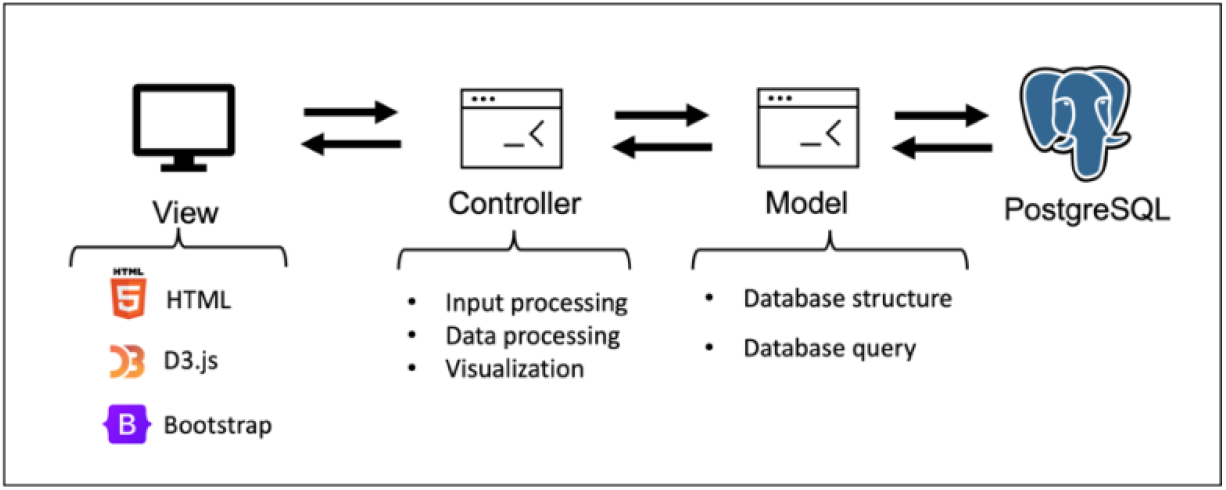
Website system design.

## Results

### Summary of the Current PubAD Publication and Analytical Results

The NLP pipeline described above has been implemented using Python libraries. We successfully constructed a gene-centric publication library that included comprehensive and summarized information related to AD research. From the data collection step, we built GSPP files for a total of 19,504 genes. With the gene-based and disease-based filtering being applied to the GSPPs in the data cleaning process, 9193 GSPPs did not contain any AD/ND related publications. The resulting remaining 10311 GSPPs were processed further for information extraction and visualization. Once the data extraction step was performed, we removed 328 GSPPs which contains no records for all three categories of information (brain region, mouse model and keywords). Finally, we have 9983 genes’ publication information from AD and other ND related publications in the current systems. For each category of information, the number of publications formed a base to provide valuable insights for AD research (Table 1).

**Table 1.**
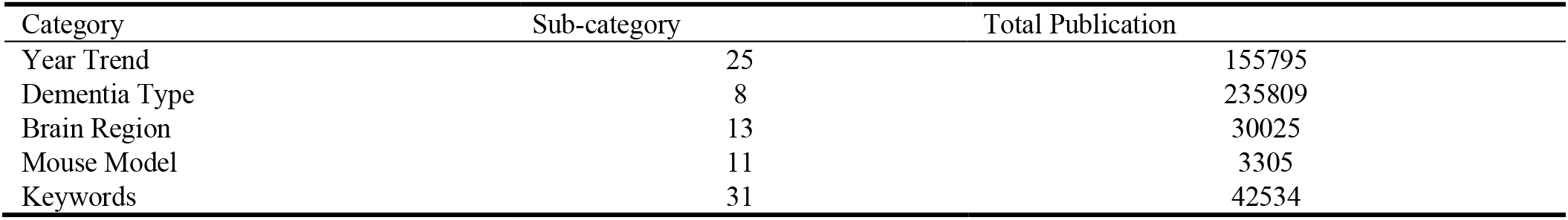
Sub-category and total publications for each information category.

To investigate the information extraction performance of our NLP pipeline, we downloaded the gene and disease annotation data from PubTator[22] database and compared the information extracted by our NLP pipeline with the annotation data from PubTator. Since PubTator does not provide ND/AD specific annotation information such as key words, brain region and mouse models, we can only compare disease and gene information. Taking Alzheimer’s disease and APOE as an example. After filtering the PubTator data, 32074 publications are annotated with Alzheimer’s Disease and APOE. In the processed PubAD data, 5395 publications are labeled by Alzheimer’s Disease and APOE. The number of overlapped publications is 4242. Then we manually examined a subset of publications annotated by AD and APOE in PubTator but not annotated in PubAD, we found the majority of these publications are recorded in PubMed Central (PMC) and PubTator used their full text data to perform annotation rather than using title and abstract, this significantly increase the number of publications annotated with AD and APOE. However, most of these non-overlapped publications are not focusing on AD and APOE research. For example, PMID30409187: is focusing on Parkinson’s Disease and APOE is only mentioned once in Methods section[23]; PMID30410670: is focusing on APOE but AD is only mentioned in Discussion section[24]. By comparing to PubTator database, we have confirmed the processed data in PubAD achieves higher specificity and provides more valuable domain specific publication information annotation such as brain region and mouse model information.

### Web Portal Development Results

On the home page of PubAD, a search box can be used to query the gene of interest by using HUGO gene symbols or ENSEMBL IDs (Figure 3). For dementia type, brain region and mouse model, a horizontal bar chart is shown on the left of the dashboard. A bar chart and a word-cloud plot are provided for visualizing the keywords information (Figure 3A). In the co-occurred genes tab, a lollipop chart is provided to demonstrate the most frequently mentioned genes of the query gene (Figure 3B). All bars of bar charts in the visualization dashboard are clickable. When clicking the bar, users can view detailed information of publications counted by clicked bar on PubMed websites. In addition, detailed publication count and subcategory information will be shown in the information box on the right when the cursor is placed on the bars (Figure 3C).

**Figure 3.**
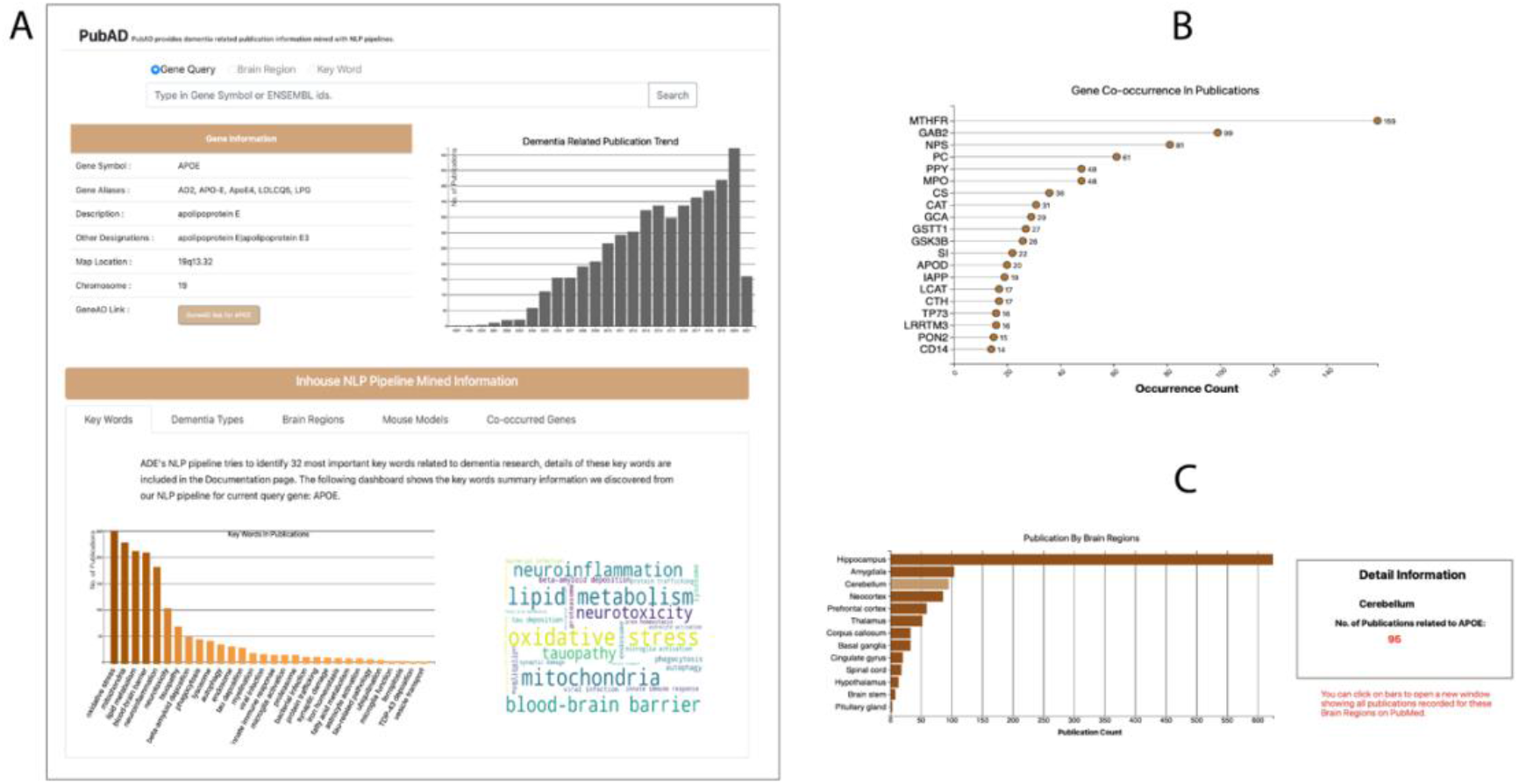
Illustrations of the PubAD portal with the sample query results of APOE. A. The general information and publication trend, along with the topics regarding the search genes and the word cloud. B: Lollipop chart for gene-gene co-occurrence in the same publication. C: The publication statistics for brain region information.

### Case Study

Using this website, we conducted a case study to show how our publication library can help researchers quickly conduct their research survey with all the summarized publication data. Figure 2A illustrated the search results of APOE, one of the most studied genes in AD or other neurodegenerative diseases. Researchers have shown immense interest in its role in ND progression, even though the underlying mechanism is still unclear [25]. Using the dataset constructed from the proposed NLP pipeline, we can conveniently check the publication trend in recent years, which is the upper right bar plot in Figure 2A. It indicates that the published studies on APOE started to rise in 2004 and hold a high increasing rate in AD/ND related fields.

The keywords occurrence information (the bottom panel of Figure 3A) is illustrated with a sorted bar plot and a word cloud plot. The visualizations help identify the most discussed keywords for the searched gene’s publications. For APOE, the top four most mentioned keywords are mitochondria, lipid metabolism, oxidative stress and blood-brain barrier. The importance of these keywords was confirmed by reviewing key scientific literature. It was recently reported that the dysfunction of mitochondria might be triggered by APOE, which plays a vital role in AD development and progression [26]. Oxidative stress’s influence on AD development has been discussed for decades. It was also recently reported that APOE4 isoform increased AD patients’ oxidative stress levels and promotes AD progression [27, 28]. Other categorical information, including the dementia types, brain regions, and mouse models, will have similar plots for the count histogram for keywords information.

The gene co-occurrence will be illustrated using a Lollipop plot (Figure 3B) which shows the number of publications that each gene (the **Y**-axis) co-occurring with the query gene (APOE). This information helps investigate related genes and pathways based on gene-gene co-occurrence in publications.

## Discussion

This study has designed and implemented an NLP pipeline to extract gene-specific ND-related publication information from the PubMed database. The processed data is stored in a structured database to enable downstream analysis. A user-friendly website has been developed for researchers to query and visualize the publication analysis results. The PubAD site has been featured and cross linked in Agora which is an important research evidence portal in AD research field[29]. To the best of our knowledge, our study is the first of its kind to categorize and summarize AD/ND related publications using an automated NLP pipeline which can provide a generalized view of gene-centric scientific research status. The processed data from the automated NLP pipeline builds a comprehensive data repository for further analysis of AD/ND related publications. For example, the processed information can be used for identifying promising drug targets by analyzing emerging research outcomes that evolve certain disease pathology, all the necessary information needed by this type of analysis has been processed in the keyword tagging step of the NLP pipeline. The processed information can also be used for the investigation of AD progression mechanisms. With the gene co-occurrence information, network analysis can be performed and mapped back to other biological networks, such as protein-protein interaction, to investigate molecule interactions [30]. Moreover, it also offers additional support for hypothesis generation and validation.

Due to the complexity of publication text mining, it is extremely difficult to conduct exhausted target information extraction in scientific literature by using an automated NLP pipeline in an unsupervised manner. Further investigations are still needed to better discriminate important information in long and complex scientific sentences. With the advance of automated annotation in biomedical literature as introduced in PubTator, more reliable training datasets will be available for building supervised information extraction pipelines. Our current NLP pipeline used expert provided domain specific key words for information extraction, to better address the publication mining related to AD/ND, more complete AD/ND ontology dictionaries are still needed to sufficiently cover all important information in AD/ND research, and it will help improve the accuracy of the keywords tagging and concept identification in AD related publications. Moreover, we have automated our NLP pipeline to run it once a month, so that we can incorporate the most recently published research outcomes, our web portal will be updated simultaneously after each run of the NLP pipeline.

## Conclusion

Our NLP pipeline and interactive web portal provide a valuable tool and data source for text mining on ND publications. The web portal is especially convenient for users without informatics skills and it will assist AD researchers in conducting preliminary examinations and learn state-of-the-art research outcomes accurately and efficiently. Our project contributes to the current AD research by providing a gene-centric publication overview on AD/ND thus will help researchers to better address the challenges in AD research.

## List of Abbreviations

AD: Alzheimer’s Disease
ND: neurodegenerative disease
NLP: natural language processing
GSPP: genespecific publication profile

## Declarations

### Ethics approval and consent to participate

Not applicable.

### Consent for publication

Not applicable.

### Availability of data and materials

The datasets used and/or analyzed during the current study are available in the PubMed Database (https://pubmed.ncbi.nlm.nih.gov), the scripts used for processing the original data are available from the corresponding author on reasonable request.

### Competing interests

The authors declare that they have no competing interests.

### Funding

This project is partially funded by the Indiana Precision Health Initiative Fund and 5U54AG065181-02 NIH funding for Alzheimer’s Disease Drug Discovery Center.

### Authors’ contributions

JL conducted the analysis and write the manuscript. JZ provided domain specific knowledge in NLP pipeline. DR, HW and JZ revised the manuscript. All authors read and approved the final manuscript.

## Acknowledgements

Not applicable.

## Notes

### Competing Interest Statement

The authors have declared no competing interest.

